# Benchmarking AI Protein Structure Predictors Reveals a Persistent Bias in Multi-State Proteins

**DOI:** 10.64898/2026.07.10.737860

**Authors:** Muhui Ye, Yu-Hong Wang, Maximilian Brogi, Jerry M. Parks, Katie M. Kuo, James C. Gumbart

## Abstract

Protein structure predictors achieve high single-state accuracy, but it remains unclear whether they can recover functionally relevant conformational ensembles or account for the presence of ligands and/or binding partners. Here, we benchmark AlphaFold3, Boltz-2, Chai-1, and BioEmu on four canonical multi-state proteins (Pf-MATE, LAO, SecA, and β_2_AR), quantifying state bias and sampling breadth against experimental reference structures. Models frequently default to a dominant state represented in the PDB; small-molecule ligands have weak or inconsistent effects, while large protein partners drive clear conformational switching between states. Multiple sequence alignment (MSA)-based approaches (AF-Cluster and random subsampling) recapitulate similar biases, indicating that this behavior is not unique to newer architectures. These results underscore current limitations for multi-state protein structure prediction and structure-guided ligand discovery.

**TOC Graphic:** 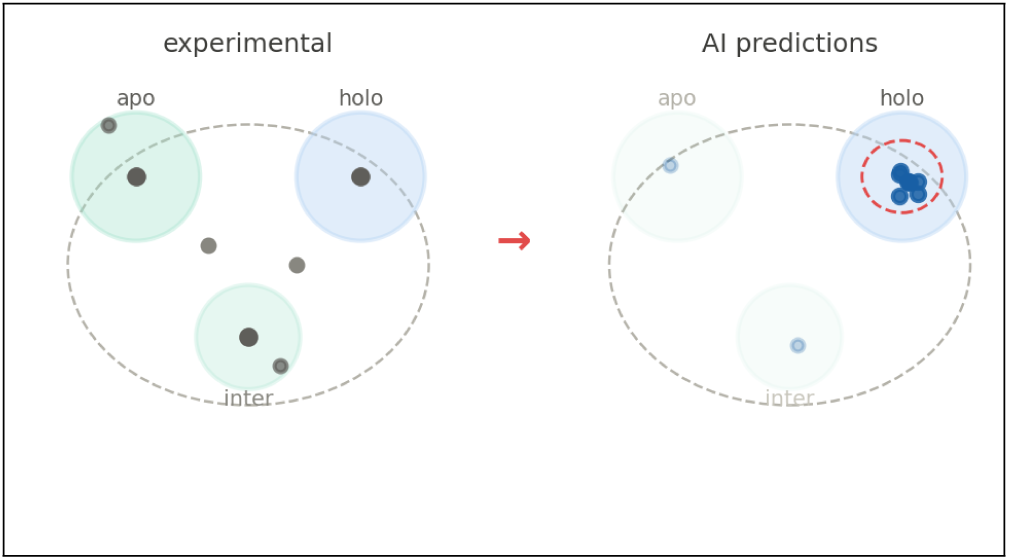

---

In 2020, AlphaFold2 revolutionized protein structure prediction by achieving near-atomic accuracy in CASP14.^1,2^ However, AlphaFold2 and related predictors often return a single dominant conformational state, limiting their ability to capture conformational heterogeneity.^3^ This limitation is important because proteins function as dynamic ensembles rather than static structures, and alternative states can control ligand recognition, allostery, transport, and catalysis.^4–6^ Prior studies have shown that AlphaFold2 predictions can be biased toward particular conformational states, in part reflecting the structural distributions represented in the Protein Data Bank (PDB) training data.^7,8^

In 2024, Abramson et al. introduced AlphaFold3, which uses a diffusion-based approach that is distinct from the architecture used in AlphaFold2. ^9^ This approach is well suited to protein structure prediction because diffusion treats the fold as a distribution over atomic coordinates, enabling AlphaFold3 to sample and rank multiple plausible structures rather than committing to a single deterministic output. It also has a reduced reliance on multiple sequence alignments (MSAs), moving closer to a truly de novo modeling paradigm with a simpler and less computationally intensive workflow. Reduced dependence on MSAs allows greater accommodation of protein sequence diversity and, consequently, conformational heterogeneity. This shift comes at the cost of increased hallucinations and unrealistic structures, particularly in intrinsically disordered or low-confidence regions.^10,11^ As a result, it remains unclear how well diffusion-based architectures, even when provided with ligand constraints, can reproduce biologically relevant protein ensembles.

Popular alternatives to AlphaFold3 employ various architectural innovations that may be useful in conformational sampling. Chai-1 (2024) is based on AlphaFold3 and includes experimental restraints, potentially improving predictions of biomolecular complexes when ligand or interaction information is available.^12^ Another predictor, Boltz-2 (2025), provides AlphaFold3-like functionality while also predicting binding affinities for ligands. ^13,14^ In addition, Boltz-2 allows user specification of protein-ligand contact restraints. Alternative protein-ligand conformations may be associated with different binding affinities, which could help identify or prioritize distinct conformational states. Lastly, BioEmu (2025) is designed to sample protein conformational ensembles using AlphaFold-derived structural representations and training data that include molecular dynamics (MD) simulations. ^15^

Given the relatively recent release of these predictors, an important question is whether they have memorized training data rather than learned generalizable molecular interaction features. While deep learning has greatly advanced protein–ligand interaction modeling,^16^ accurately capturing partner-driven conformational changes remains challenging, as protein–protein interfaces are generally more complex and flexible than small-molecule binding sites. Studies have shown that AlphaFold3 can differentiate some true binders from non-binders, but also misses many binders.^17,18^ These studies further showed that performance depends strongly on similarity to training data. Despite architectural advances beyond AlphaFold2, training-set bias in AlphaFold3 has repeatedly been found to persist, especially when modeling alternative conformations.^19–21^ Although MD simulations can expand training data beyond static PDB structures, they do not necessarily eliminate training-set bias, and tools such as BioEmu may still generate predictions with spurious flexibility. ^22,23^ Boltz-2, which builds on Boltz-1’s demonstrated strength in diverse ligand-protein multimer structure prediction, also included MD data during training. However, it may fall short in exploiting these data for conformational modeling because its emphasis on affinity prediction remains.^13,24^ Although Chai-1 shows competitive accuracy relative to AlphaFold2 and Boltz-1 in recovering known conformations and representing transition trajectories, recent benchmarks suggest that Chai-1, like AlphaFold3 and Boltz-2, remains unable to recapitulate common conformational rearrangements.^18^

We are therefore motivated to test whether these predictors can recover experimentally observed conformational states, and whether ligand or binding-partner specification improves conformational sampling beyond biases inherited from training data. Here, we systematically evaluate AlphaFold3, Boltz-2, Chai-1, and BioEmu across four well-characterized proteins with distinct experimentally resolved conformations: PfMATE, a multidrug transporter with inward- and outward-facing apo conformations;^25^ LAO, a periplasmic binding protein with a ligand-induced open-to-closed transition;^26,27^ SecA, an ATPase with nucleotide- and partner-dependent conformational states;^28,29^ and β_2_-adrenergic receptor (β_2_AR), a GPCR with active and inactive conformations.^30^ These proteins represent biological functions in which conformational switching is directly responsible for activity, regulation, ligand recognition, or a combination thereof, making them useful benchmarks for evaluating conformational sampling in proteins that undergo similar transitions.

## Test case 1: PfMATE

The multidrug and toxic compound extrusion (MATE) transporter family enables multidrug resistance in bacterial pathogens as well as cancer cells by using electrochemical ion gradients (typically H^+^ or Na^+^) to export xenobiotics and cytotoxic compounds across the cell membrane.^31,32^ The proton-coupled conformational transition between outward-facing (OF) and inward-facing (IF) states involves a rocker-switch mechanism in which transmembrane (TM) helices undergo coordinated movements, forming the characteristic inverted V-shape of the IF conformation.^25,33^ PfMATE is a MATE transporter from the hyper-thermophilic archaeon *Pyrococcus furiosus*.^25^ PfMATE is a well-characterized model system for studying MATE transporter conformational dynamics, with both inward-facing (PDB: 6FHZ) and outward-facing (PDB: 6GWH) structures experimentally resolved.^34^

Chai-1 and Boltz-2 showed a similar bias toward the OF state, with nearly identical performance (RMSD to OF: 0.79 ± 0.09 Å and 0.74 ± 0.11 Å, respectively; RMSD to IF: 5.10 ± 0.21 Å and 4.93 ± 0.26 Å, respectively). The low standard deviations indicate minimal conformational sampling, whereas AlphaFold3 was even less (RMSD to OF: 0.57 ± 0.03 Å; RMSD to IF: 5.42 ± 0.05 Å). Thus, all three tools converged on tightly clustered predictions around the OF state.

BioEmu displayed substantially broader conformational sampling compared to the other tools, with higher standard deviations (RMSD to OF: 1.22 ± 0.40 Å; RMSD to IF: 3.56 ± 0.90 Å). The predictions are more dispersed in conformational space, demonstrating that its MD-based training strategy enables greater exploration (Figure 1). Notably, some BioEmu frames approached intermediate RMSD values (2-3 Å to both reference structures), indicating partial sampling of conformational states between IF and OF. However, despite this broader distribution, nearly all BioEmu predictions remained closer to the OF state, and only a single prediction was closer to the IF state.

**Figure 1:**
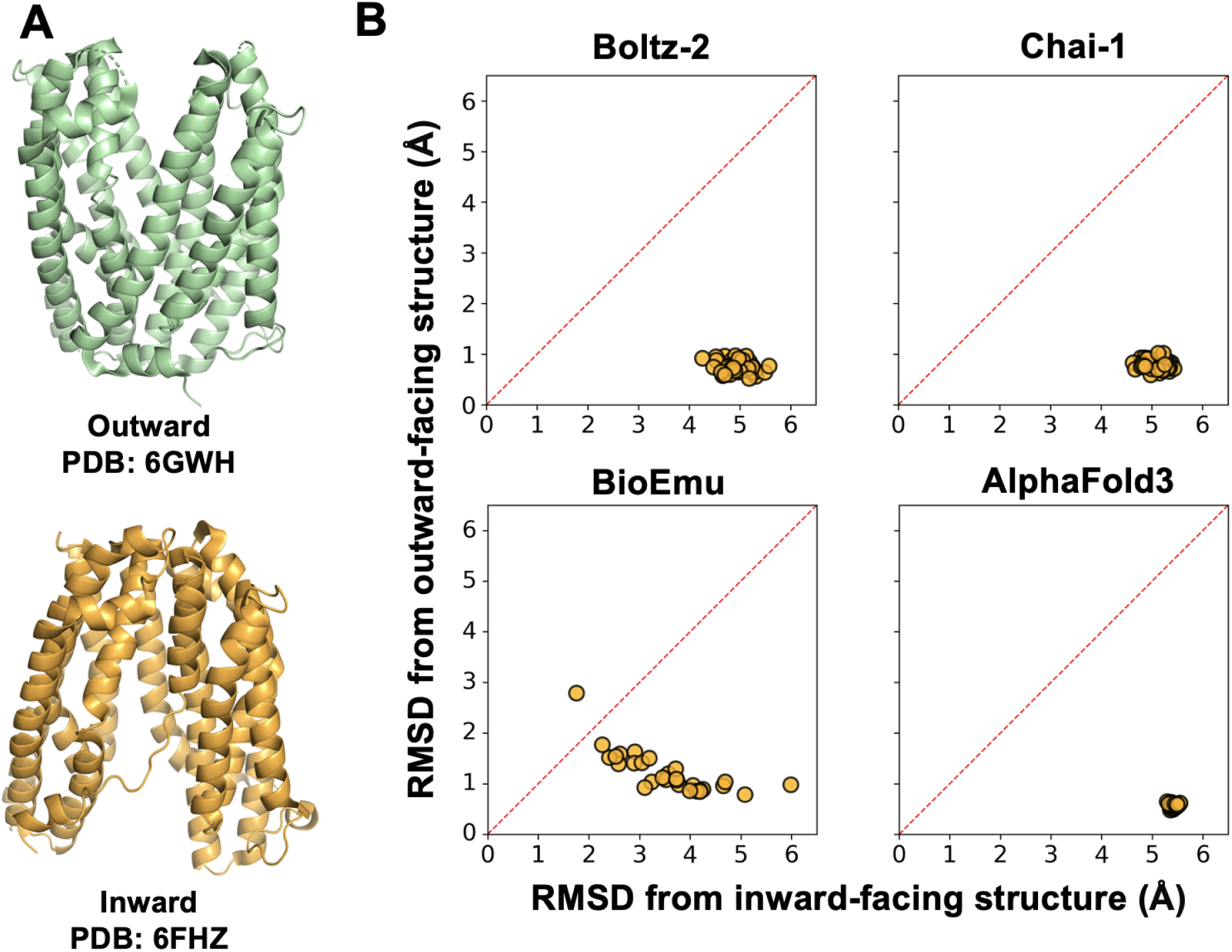
Conformational analysis of PfMATE predictions. (A) Reference structures showing the OF state (PDB: 6GWH, green) and IF state (PDB: 6FHZ, yellow). (B) RMSD of predicted structures from Boltz-2, Chai-1, BioEmu, and AlphaFold3 compared to IF and OF reference structures. Points below the diagonal line are closer to the OF state, while those above are closer to the IF state.

Except BioEmu, which generated one structure 1.8 Å away, no tool successfully predicted structures resembling the IF reference state; most RMSD values to the IF state exceeded 3.5 Å (Figure 1B). This bias occurred despite the IF-state structure being released in 2019 and therefore present in the training data for all models tested. Existing structures of Pf-MATE predominantly adopt OF conformations. Of the eleven PfMATE structures deposited in the PDB, only one (6FHZ) captures the IF state, creating a 10:1 ratio that may contribute to the observed prediction bias (Table S1). All models converged to majority (or exclusive) OF-state predictions regardless of their architecture. Boltz-2 and Chai-1 showed nearly identical performance, while BioEmu had broader sampling yet failed to fully reproduce the IF conformation. AlphaFold3 produced the tightest clustering around the OF state.

## Test case 2: LAO binding protein

The lysine/arginine/ornithine-binding protein (LAO) from *Salmonella typhimurium* is a periplasmic binding protein that undergoes large-scale domain rearrangements upon binding its cognate amino acid ligands (lysine, arginine, or ornithine).^26^ The protein transitions between an open (apo) state (PDB: 2LAO)^26^ and a closed (holo) state (PDB: 1LAF),^35^ in which the inter-lobe closure is enabled by an ∼52*^◦^* rotation of a backbone torsion angle in the connecting strand.^26^ Having a “Venus-flytrap” mechanism characteristic of the periplasmic binding protein superfamily, LAO initiates signal transduction in bacterial chemotaxis and nutrient uptake systems.^36^

AlphaFold3 and Chai-1 showed similar behavior, with a strong bias toward the closed holo conformation across both prediction conditions (apo and with the ligand arginine). For apo predictions, both tools converged tightly to the holo state (AlphaFold3: RMSD to holo = 0.20 ± 0.01 Å, RMSD to apo = 4.63 ± 0.03 Å; Chai-1: RMSD to holo = 0.29 ± 0.03 Å, RMSD to apo = 4.60 ± 0.06 Å). The holo predictions showed nearly identical performance (AlphaFold3: RMSD to holo = 0.22 ± 0.02 Å; Chai-1: RMSD to holo = 0.24 ± 0.02 Å). The very low standard deviations (0.01-0.06 Å) across all conditions indicate minimal conformational sampling, with predictions tightly clustered around the holo state regardless of input conditions.

Boltz-2 demonstrated a critical distinction between its prediction conditions. Although holo predictions converged tightly on the holo state (RMSD to holo = 0.27 ± 0.03 Å, RMSD to apo = 4.62 ± 0.04 Å) with high model confidence (pTM 0.929–0.952), apo predictions exhibited substantially broader conformational variance (RMSD to holo = 0.58 ± 0.75 Å, RMSD to apo = 4.48 ± 0.74 Å) and markedly lower confidence (pTM 0.729–0.856, Table S4). Among 50 apo predictions, two structures approached the open apo conformation, while the remaining 48 converged on the holo state (Figure 2). These two successful predictions represent the only instances across all tested models in which apo structure predictions approached the correct open conformation.

**Figure 2:**
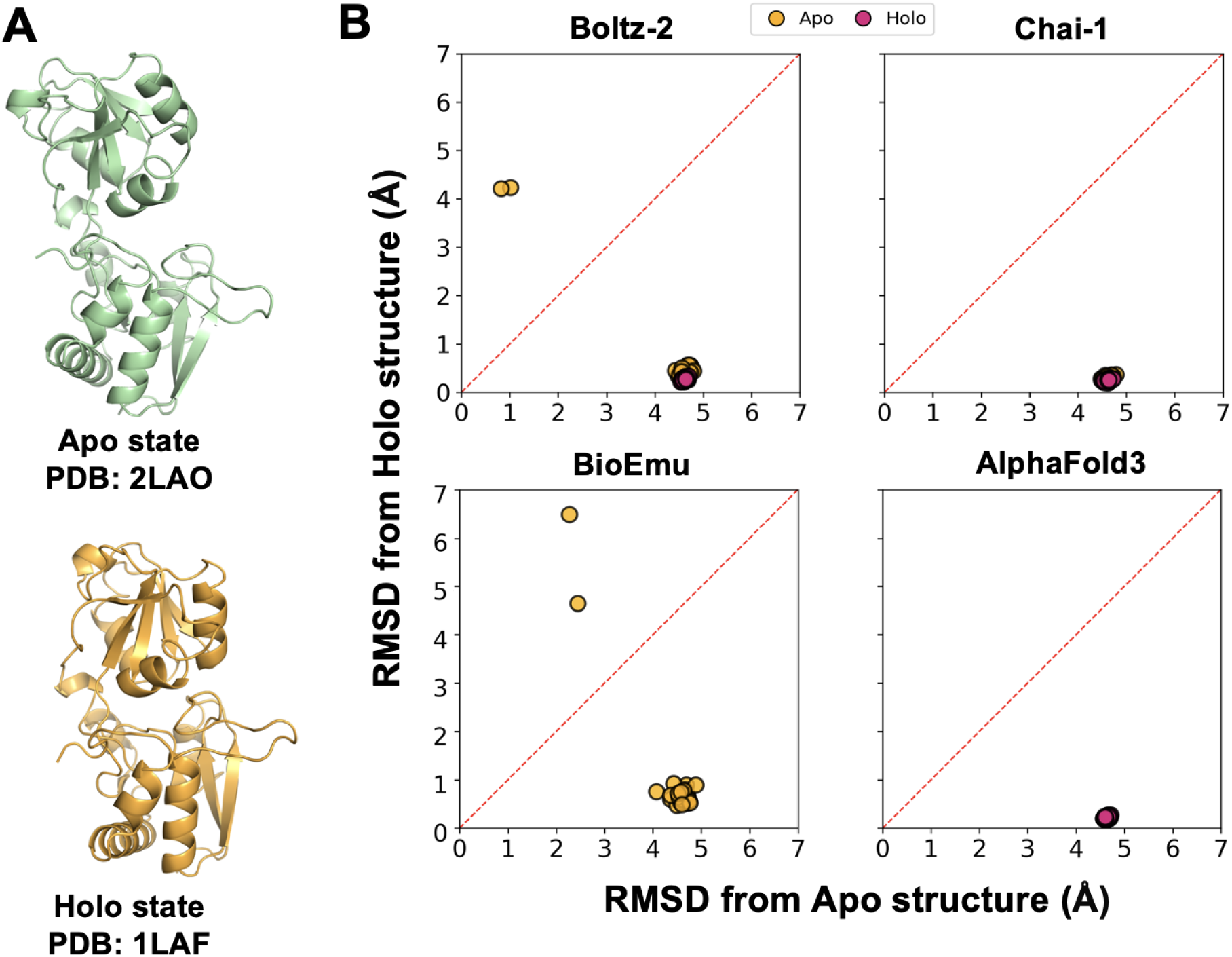
Conformational analysis of LAO predictions. (A) Reference structures showing the apo (open, PDB: 2LAO, green) and holo (closed, PDB: 1LAF, orange) conformations. (B) RMSD of predicted structures from Boltz-2, Chai-1, BioEmu, and AlphaFold3 compared to apo and holo reference structures. Data are shown as circles in two input conditions: apo (orange) and holo (pink). Points below the diagonal line are closer to the holo state, while those above are closer to the apo state.

BioEmu demonstrated broader conformational sampling with higher standard deviations (RMSD to apo: 4.42 ± 0.63 Å; RMSD to holo: 1.04 ± 1.36 Å). Just as for PfMATE, BioEmu predictions are more dispersed across conformational space, with some frames displaying intermediate RMSD values to both reference structures (Figure 2). However, despite this broader sampling, BioEmu did not predict the open apo state. All BioEmu frames remained biased toward the holo conformation, with even the most apo-like frames (minimum RMSD to apo: 2.27 Å) falling short of the experimental open structure.

For LAO, we provided explicit ligand information to AlphaFold3, Chai-1, and Boltz-2 (arginine in holo runs; no ligand in apo runs). Despite this specification, the vast majority of apo predictions converged to the closed holo conformation. AlphaFold3 and Chai-1 produced nearly identical apo and holo predictions, whereas Boltz-2 showed greater apo-state variability but remained predominantly holo-like.

## Test case 3: SecA

SecA is an ATPase motor protein that drives bacterial preprotein translocation through the SecYEG channel.^37,38^ Its structure includes a helicase-like core composed of two nucleotide-binding domains (NBD1 and NBD2), a helical wing domain (HWD), and a helical scaffold domain (HSD). Crucially, the preprotein binding domain (PBD), NBD2, and HSD form a functional clamp that captures polypeptides above the SecY pore. ^39–41^

In its apo state, SecA exists as a dynamic ensemble. Single-molecule FRET shows the PBD is highly mobile, sampling multimodal positions relative to the HWD and NBD2.^42^ This heterogeneity is seen in experimental structures such as the wide-open state (PDB: 1M74, ref.^28^), where the PBD resides near the HWD, and the intermediate-open state (PDB: 1TF2, ref.^43^). ATP binding shifts this equilibrium toward the wide-open conformation by drawing the PBD closer to the HWD.^42^

Upon docking onto SecYEG with ADP bound, SecA undergoes a large conformational change in which the PBD rotates toward NBD2, closing the clamp around the translocating polypeptide.^39,42^ Recent cryo-EM structures of the SecA-SecYEG complex, e.g., PDB: 8YAS,^44^ provide a high-resolution view of this engaged state. During active translocation, SecA is believed to employ a push-and-slide mechanism: ATP binding drives the two-helix finger to push the polypeptide into the channel, while subsequent hydrolysis is coupled to clamp closure that limits backsliding of the substrate. ^38,40^

Because SecA conformational changes are primarily localized to the rotation of the PBD domain rather than global structural rearrangements, RMSD of the entire protein is insufficient to distinguish between open and closed clamp states. We therefore quantified the conformational ensemble of each prediction using two domain-level metrics: the PBD–HWD distance and the PBD–hinge–NBD2 angle, which together capture the extent of clamp opening. Reference values were taken from experimental structures representing three key states: wide-open (1M74; distance = 26.7 Å, angle = 135.0*^◦^*), open (1TF2; 39.3 Å, 106.0*^◦^*), and closed (8YAS; 49.9 Å, 76.1*^◦^*).

In the apo condition, AlphaFold3 predictions cluster predominantly around the wide-open state (29.4 ± 5.4 Å, 126.9 ± 10.0*^◦^*), close to 1M74 (26.7 Å, 135.0*^◦^*), with a small fraction (6 out of 50) approaching the open conformation (Figure 3). Chai-1 and Boltz-2, however, fail to sample the intermediate open state: their predictions are bimodally distributed between closed and wide-open conformations, with no structures near the open reference state, indicating that these models cannot reproduce the conformational heterogeneity characteristic of apo SecA. BioEmu presents a distinct behavior, sampling a broad continuous distribution from wide-open to closed (45.1 ± 4.3 Å), suggesting that it captures a more complete conformational landscape, although one that is unphysiologically biased towards the closed state.

**Figure 3:**
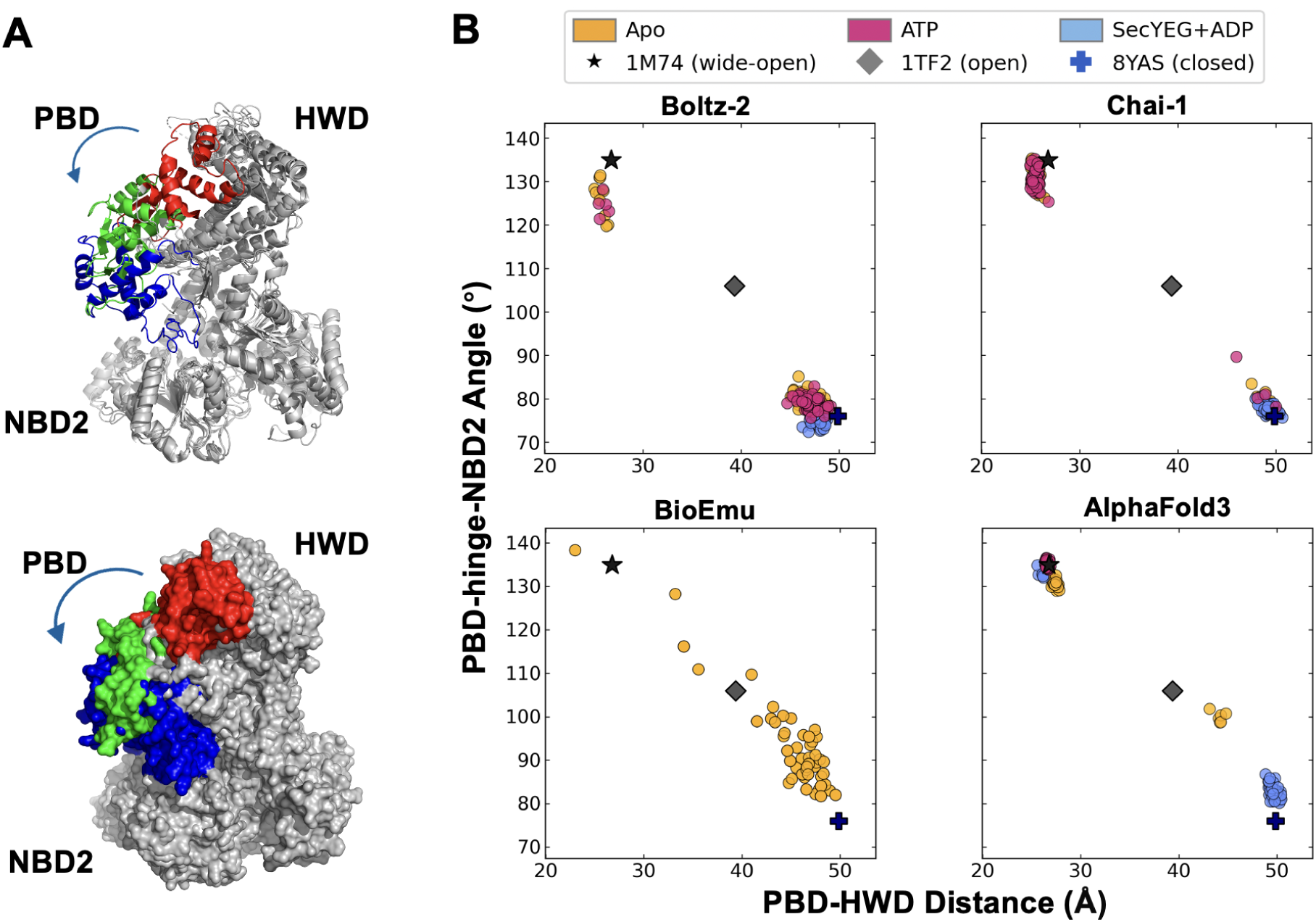
Conformational analysis of SecA predictions. (A) Reference structures of SecA in three conformational states shown in cartoon (top) and surface (bottom) representations: wide-open (1M74, red), open (1TF2, green), and closed (8YAS, blue). The arrow indicates the rotational movement of the PBD domain. (B) PBD–HWD distance and PBD–hinge–NBD2 angle of predicted structures from Boltz-2, Chai-1, BioEmu, and AlphaFold3 under three conditions: apo (orange), with ATP (pink), and with SecYEG and ADP (blue). Reference structures are indicated on each plot as a black star (1M74, wide-open), a gray diamond (1TF2, open), and a blue cross (8YAS, closed).

With ATP included as a ligand, a wide-open conformation is expected. AlphaFold3 correctly predicts this state, collapsing to a tight cluster near 1M74 (26.6 ± 0.1 Å). Chai-1 and Boltz-2 again fail to capture the expected response, with both models producing a mixture of closed-like and wide-open predictions rather than converging on the wide-open state, mirroring the same bimodal failure observed in the apo condition.

Upon providing SecYEG and ADP, clamp closure is expected, as exemplified by the structure in PDB 8YAS. Chai-1 and Boltz-2 both correctly converge on the closed conformation, with mean distances of 49.3 ± 0.5 Å and 47.8 ± 0.5 Å, closely matching 8YAS (49.9 Å) and exhibiting low variance. AlphaFold3, however, fails in this condition (46.9 ± 7.7 Å): a substantial fraction of predictions remains in the wide-open state, reflecting an inconsistent response to the binding of SecYEG and ADP. Model confidence was generally lower under SecYEG+ADP conditions (pLDDT 72–81) compared to apo predictions (pLDDT 76–86, Table S4), consistent with the increased structural complexity of the multiprotein complex.

## Test case 4: β_2_AR

The β_2_-adrenergic receptor (β_2_AR) is a G protein-coupled receptor (GPCR) that is responsible for regulating physiological responses to agonists such as epinephrine and nore-pinephrine, facilitating cardiovascular function, smooth muscle relaxation, and metabolic regulation.^45^ β_2_AR is well characterized and widely conserved across mammals and birds.^46^ Agonist binding alone stabilizes intermediate receptor conformations but is generally insufficient to induce the fully active state; the binding pocket exhibits only around a 1-Å shift upon agonist binding.^47^ Full activation requires engagement with the heterotrimeric G protein, which allosterically stabilizes the outward displacement of TM6 at around 14 Å.^47–49^ This layered activation mechanism is challenging for structure prediction models, as it spans a wide conformational space. Nevertheless, defining input conditions by their bound ligands and partners allows an opportunity to investigate whether these can guide models toward the desired conformational states.

The largest conformational changes of the GPCR stem from the TM helices during its activation.^50^ Among the four input conditions (apo [inactive], agonist [inactive], agonist-Gαβγ [active], and agonist-Gαβγ-GTP [active]), the active and inactive states are the main conformations of interest as they are the most distinct. Figure 4 demonstrates that all baseline predictions (yellow dots) without user-specified ligands show weak bias toward the inactive conformation with the exception of Chai-1, which samples both states (average RMSD to the inactive structure: Boltz-2, 2.12 ± 0.33 Å; Chai-1, 2.91 ± 1.00 Å; BioEmu, 2.15 ± 0.46 Å; AlphaFold3, 1.65 ± 0.05 Å). Chai-1 identifies clusters within both conformations without any constraints. When provided with the partners and ligands required for activation of β_2_AR (i.e., agonist and G protein), all predictions clustered tightly toward the active conformation (average RMSD: Boltz-2, 1.68 ± 0.33 Å; Chai-1, 1.02 ± 0.07 Å; AlphaFold3, 0.74 ± 0.02 Å).

**Figure 4:**
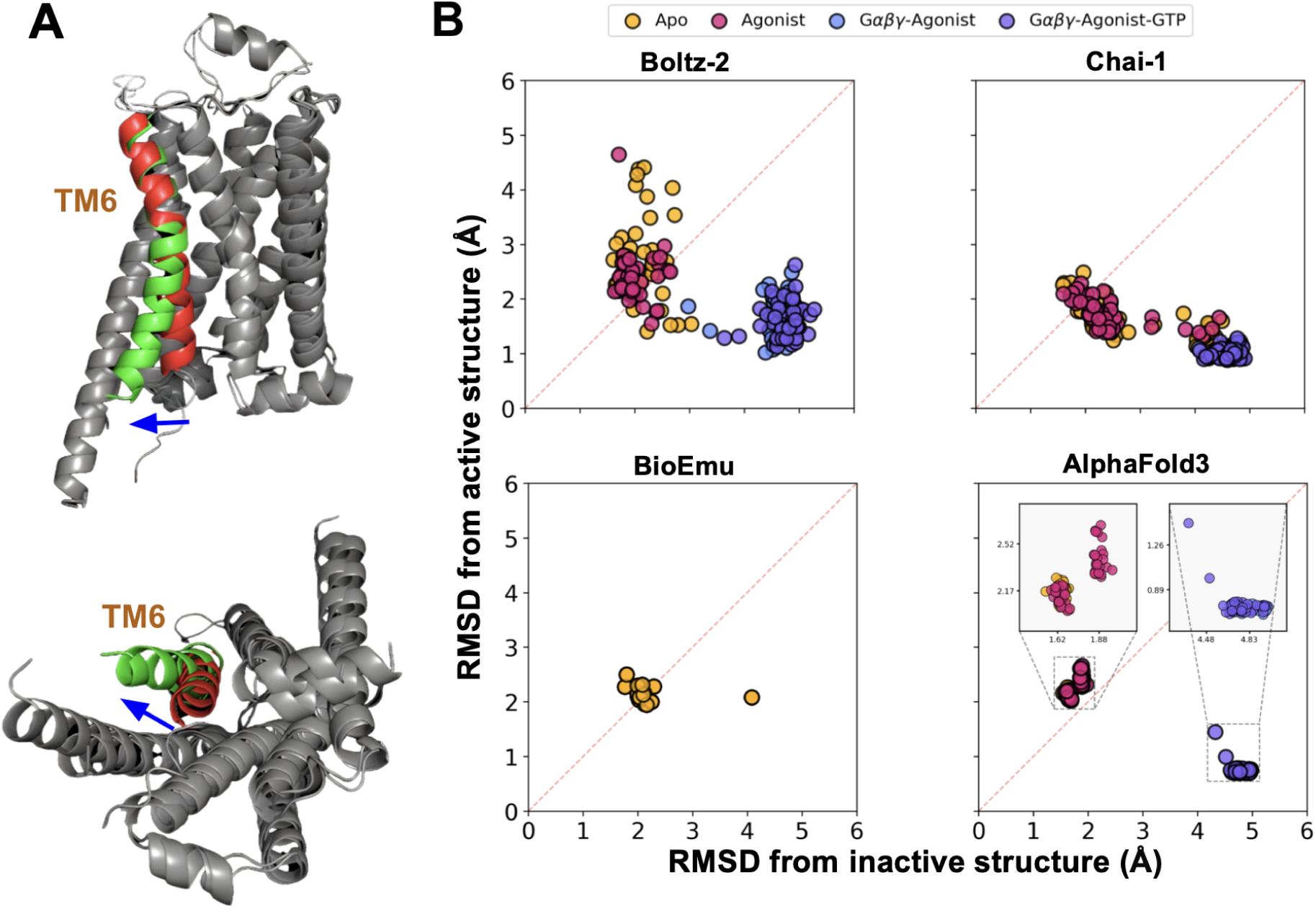
Conformational analysis of β_2_AR predictions. (A) Aligned reference structures highlighting the shift of TM6, from 9CHU (red, inactive, agonist-bound structure) to 8GEG (green, active, agonist-Gαβγ bound), where the top and bottom images display the side and cytoplasmic views, respectively. (B) RMSD of predicted structures from Boltz-2, Chai-1, BioEmu, and AlphaFold3 compared to inactive (9CHU) and active (8GEG) reference structures. Data are shown as circles in four input conditions: apo (orange), agonist (pink), Gαβγ-agonist (blue), and Gαβγ-agonist-GTP (green). Points below the diagonal line are closer to the active state, while those above are closer to the inactive state. The small cluster generated by BioEmu shows 20 predicted structures, and AlphaFold3 predictions of Gαβγ-Agonist overlap with Gαβγ-Agonist-GTP.

Notably, predictions with the G protein partner (blue, purple) achieved substantially lower RMSD to the active reference structure than baseline apo and agonist-only predictions (yellow, pink) did to the inactive reference. Chai-1 and AlphaFold3 produced particularly tight clusters for the active state. In contrast, baseline predictions remained intermediate between the two reference structures rather than cleanly adopting the inactive conformation (Figure 4). These results indicate that Boltz-2, Chai-1, and AlphaFold3 may favor intermediate or weakly inactive-like states until provided with the appropriate partner and ligand configuration. Even without ligands or partners, the predictors did not clearly recover inactive structures, whereas partner-containing inputs captured the active conformation with clear separation from the inactive reference. One BioEmu prediction and a few apo and agonist-only Chai-1 predictions also fell near the active-state clusters formed under activation-associated input conditions.

## Alternative conformational sampling strategies

The sampling limitations observed across PfMATE, LAO, SecA, and β_2_AR can potentially be addressed by using AlphaFold2, which can be forced to generate alternative conformations through manipulation of the inputs. For example, reducing the depth of MSAs via subsampling can yield alternative transporter, receptor, and fold-switching states.^8^ MSA clustering, especially AF-Cluster, goes further by separating conflicting evolutionary signals and has been experimentally validated on metamorphic proteins, making it one of the strongest demonstrations that AlphaFold2 can be redirected toward functionally relevant alternative substates.^51^ AFsample2T, which uses MSA column masking to reduce coevolutionary signals, has also been shown to produce greater structural heterogeneity than standard approaches.^52^ Alternatively, providing state-specific templates can steer proteins such as GPCRs toward active or inactive conformations when the target state is known a priori, ^3^ which is also a capability now shared by AlphaFold3, Boltz-2, and Chai-1 as well.^9,12,14^

Additional methods seek to increase ensemble diversity by altering either the MSA itself or inference stochasticity. SPEACH AF uses targeted MSA mutagenesis to redirect AlphaFold2 toward alternative conformers, ^53^ whereas ColabFold facilitates practical sampling through shallow MSAs, dropout, and multiple random seeds.^54^ Retrained models such as Cfold seek to determine whether unobserved alternative conformational states can be predicted without simple training-set recall.^55^ Physics-augmented AF2-RAVE workflows extend this approach by using AlphaFold2 structures as seeds for subsequent enhanced-sampling MD simulations and Boltzmann-like ranking.^56,57^ Another physics-based approach, DeepPath, complements such efforts by using force-field-guided active learning to generate transition pathways between protein conformational states.^58^

Confidence metrics such as pLDDT and PAE are useful for local geometry and uncertainty, but they are not reliable selectors of biologically or physically meaningful alternative states. Benchmark studies explicitly warn that apparent success can be inflated by training-set overlap or memorization.^55,59^ Likewise, even successful shallow-MSA or AF-Cluster workflows generally recover feasible substates rather than full free-energy landscapes. CASP15 and CASP16 showed that detailed multi-state prediction remains substantially weaker than single-state prediction, especially without templates and for larger assemblies or nucleic-acid-containing systems.^60–62^ Deep-learning approaches now offer useful heuristics for ensemble generation, but these ensembles do not necessarily improve downstream docking and cannot yet provide thermodynamically consistent conformational landscapes.^63,64^

To provide a baseline comparison using an AlphaFold2-based MSA manipulation approach distinct from the newer architectures evaluated above, we applied AF-Cluster^51^ to all four target proteins. AF-Cluster uses DBSCAN to partition the MSA into sequence-similarity clusters, each reflecting a distinct evolutionary subfamily that may encode a different conformational state; AlphaFold2 is then run independently on each cluster’s sub-MSA. As controls, we also generated uniformly random sub-MSAs of depth 10 (U10) and 100 (U100) to test whether evolutionary structure in the clusters, as opposed to simple MSA depth reduction, drives any observed conformational shift. As shown in Figure 5, AF-Cluster and random subsampling (U10, U100) produced similar conformational distributions for Pf-MATE, LAO, and β_2_AR, with predictions consistently favoring holo-like or single dominant states over the alternative reference conformation. SecA was a notable exception: cluster, U10, and U100 predictions were broadly scattered and did not converge to any of the three reference states (1M74, 1TF2, 8YAS), reflecting the low sequence diversity of the SecA MSA, which yielded only four DBSCAN clusters of size four to five sequences each. These results indicate that MSA-level manipulation alone, whether through evolutionary clustering or random subsampling, is largely insufficient to overcome the systematic conformational bias observed across the deep learning tools evaluated in this study.

**Figure 5:**
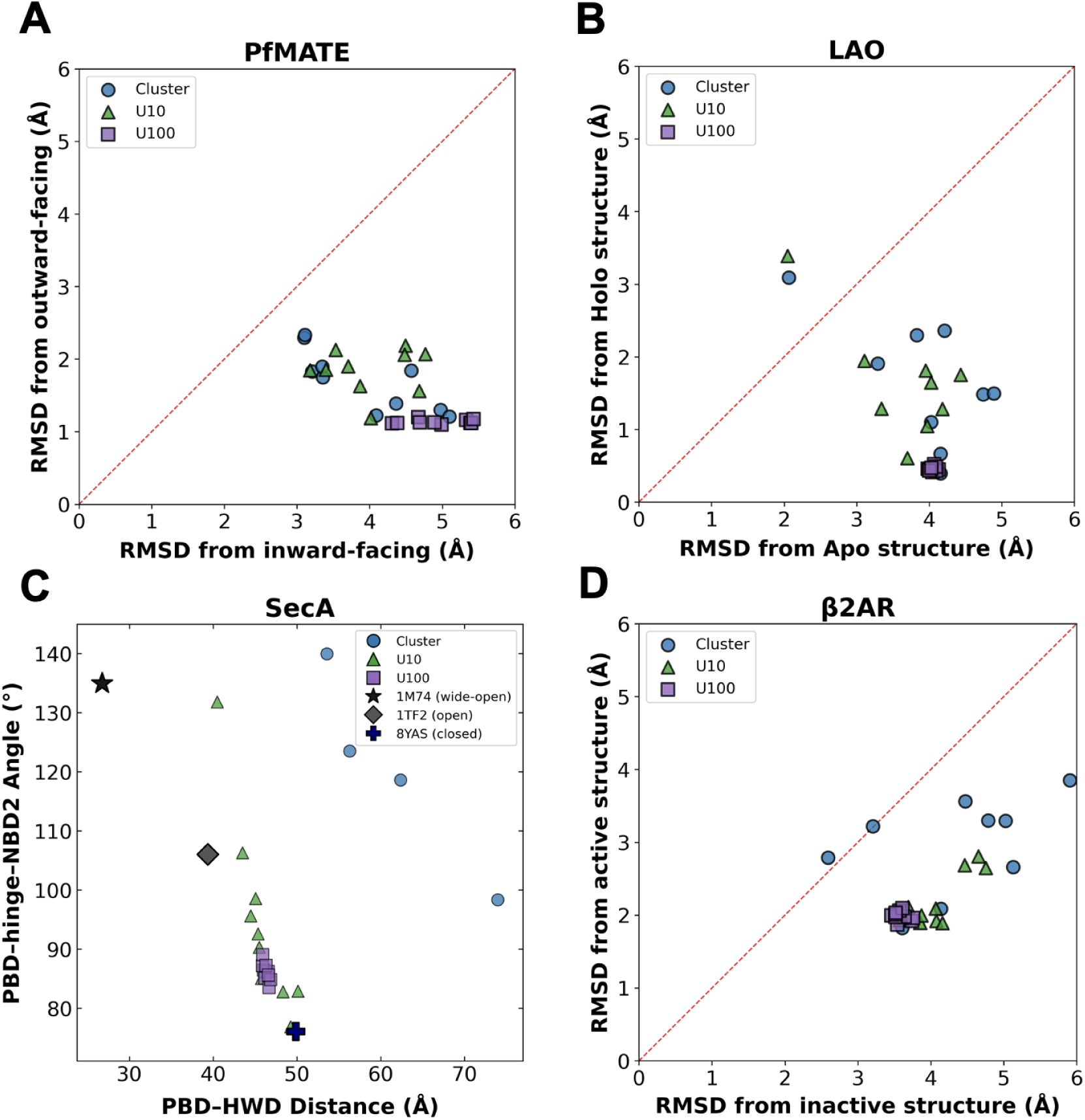
AF-Cluster and MSA subsampling results for four target proteins. Predicted structures from DBSCAN-derived MSA clusters (blue circles), uniformly subsampled MSAs of depth 10 (U10, green triangles), and depth 100 (U100, purple squares) are plotted against paired reference conformations for (A) PfMATE, (B) LAO, and (D) β_2_AR, and as PBD–HWD distance versus PBD–hinge–NBD2 angle for (C) SecA, with reference experimental structures shown as black symbols (star: 1M74 wide-open; diamond: 1TF2 open; cross: 8YAS closed). All three sampling strategies tend to favor dominant or holo-like conformations, mirroring the biases observed in the newer tools evaluated in this study.

## Conclusions

Across PfMATE, LAO, SecA, and β_2_AR, modern predictors (AlphaFold3, Boltz-2, Chai-1, BioEmu) show state-dependent success but repeatedly collapse toward a single dominant basin, consistent with training-distribution bias. As seen for AlphaFold2,^65^ AlphaFold3, Boltz-2, and Chai-1 often form compact clusters, and even without templates or MSAs, predictions frequently regress toward the dominant represented state in the PDB.^66^ Although BioEmu samples broader conformational distributions than the other predictors, its ensembles remain biased toward dominant states and generally fail to recover the alternative experimental conformations, complicating functional interpretation. Consistent with this, recent studies indicate that BioEmu can reproduce basic dynamical signatures (e.g., flexibility and correlated motions) but fails to generate energetically weighted Boltzmann ensembles and provides limited improvement for ensemble docking.^67,68^ In systems with multiple thermodynamically accessible apo conformations, predictors often recognize only one end of the ensemble (PfMATE; Figure 1) or yield weak, skewed sampling (SecA; Figure 3). Notably, the same collapse toward dominant or holo-like states observed across the newer predictors was also recapitulated by MSA-based approaches, indicating that MSA manipulation alone is insufficient to overcome the conformational bias observed here (Figure 5). This pattern is consistent with the unequal distribution of PDB structures across conformational states for all four proteins (Table S1).

Small-molecule ligand conditioning is generally weak and sometimes idiosyncratic. For LAO, including arginine did not induce alternate conformations, and apo predictions largely remained holo-like (Figure 2). This observation aligns with a recent analysis of 82 enzymes showing that apo/holo predictions from AlphaFold3 are driven primarily by the apo/holo ratio in the PDB training set rather than by ligand identity, with non-binding ligands inducing domain motions similar to true triggers.^69^ For SecA, ATP produced the expected wide-open shift for AlphaFold3 but not for Boltz-2 or Chai-1 (Figure 3), indicating that small-molecule inputs do not reliably steer predictions toward the expected functional state. Together with prior evidence of limited multimer performance in positioning small-molecule ligands, ^70^ these results suggest that current predictors do not consistently encode small-molecule-driven induced-fit mechanisms at the level needed for state control. Studies have also reported that Boltz-2 generalizability degrades for proteins with limited training coverage;^71,72^ consistent with this finding, even for canonical proteins in our benchmark, Boltz-2 does not reliably sample alternative conformational states without additional binding/affinity information.

In contrast, protein-binding partners provide a stronger and more reproducible control signal, consistent with models responding more robustly to protein–protein interface geometry/contact density than to small-molecule chemistry. For β_2_AR, providing the agonist together with the heterotrimeric G protein shifted predictions toward the expected active conformation across predictors. For SecA, partner-driven effects were strongly model-dependent: Chai-1 and Boltz-2 succeeded only with SecYEG and ADP bound, whereas AlphaFold3 captured open and wide-open states under apo and ATP-bound conditions. This complementary failure pattern suggests that each model responds to different structural cues, with partner success likely reflecting sensitivity to the large translocase interface, rather than consistently encoding the mechanochemical coupling between nucleotide state and translocase engagement that drives the conformational cycle of SecA.

## Methods

All predictions were generated de novo for four multi-state proteins (PfMATE, LAO, SecA, and β_2_AR) using AlphaFold3, Chai-1, Boltz-2, and BioEmu.^9,12,14,15^ Protein sequences and annotations were obtained from UniProt,^73^ and experimental reference structures were taken from the Protein Data Bank.^74^ Small-molecule ligands (L-arginine, ATP/ADP, nore-pinephrine, and GTP) were provided as canonical SMILES strings where supported, and protein binding partners (SecYEG; heterotrimeric G protein) were provided as full-length sequences; no distance restraints, templates, or MSAs were used. Calculations were performed on NVIDIA A100 GPUs. As a baseline distinct from the newer architectures, we additionally applied AF-Cluster,^51,75^ which partitions the MSA into sequence-similarity clusters via DB-SCAN for input to AlphaFold2, together with uniformly random MSA subsampling controls of depth 10 and 100 (U10, U100). Reference structures were preprocessed to retain only the target protein chains and remove non-protein components. Conformational similarity was quantified primarily by C*α* RMSD in PyMOL using CEAlign and sequence-based align,^76^ and conformational preferences were visualized by paired-reference RMSD scatter plots. For SecA, where three reference states and large domain motions can make RMSD ambiguous, we additionally classified states using domain-level geometric descriptors (PBD–HWD distance and a PBD–hinge–NBD2 angle). TM-scores were computed with TM-align to confirm global fold preservation.^77,78^ Complete protocols, parameters, and sampling details are provided in the Supporting Information.

## Acknowledgments

This work was supported by the National Institutes of Health (R01-GM148586) and the Georgia Research Alliance. Structure predictions were run on the Phoenix research computing cluster, which is managed by the Partnership for an Advanced Computing Environment at Georgia Tech. J.M.P. was supported by the Laboratory Directed Research and Development program at Oak Ridge National Laboratory, which is managed by UT-Battelle, LLC under contract DE-AC05-00OR22725 for the US Department of Energy.

## Data availability

Structure predictions were generated using AlphaFold3 v3.0.1, ^9^ Chai-1,^12^ Boltz-2,^14^ and BioEmu.^15^ Predictions were generated using local installations through the corresponding repositories with default parameters unless otherwise noted: AlphaFold3 (https://github.com/google-deepmind/alphafold3), Chai-1 (https://github.com/chaidiscovery/chai-lab), Boltz-2 (https://github.com/jwohlwend/boltz), and BioEmu (https://github.com/microsoft/bioemu). All outputs are available at https://doi.org/10.5281/zenodo.20077837.

## References

(1) Jumper, J.; Evans, R.; Pritzel, A.; Green, T.; Figurnov, M.; Ronneberger, O.; Tunyasuvunakool, K.; Bates, R.; Žídek, A.; Potapenko, A.; Bridgland, A.; Meyer, C.; Kohl, S. A. A.; Ballard, A. J.; Cowie, A.; Romera-Paredes, B.; Nikolov, S.; Jain, R.; Adler, J.; Back, T.; Petersen, S.; Reiman, D.; Clancy, E.; Zielinski, M.; Steinegger, M.; Pacholska, M.; Berghammer, T.; Bodenstein, S.; Silver, D.; Vinyals, O.; Senior, A. W.; Kavukcuoglu, K.; Kohli, P.; Hassabis, D. Highly accurate protein structure prediction with AlphaFold. Nature 2021, 596, 583–589.

(2) Kryshtafovych, A.; Schwede, T.; Topf, M.; Fidelis, K.; Moult, J. Critical assessment of methods of protein structure prediction (CASP)–Round XIV. Proteins: Struct., Funct., Bioinf. 2021, 89, 1607–1617.

(3) Heo, L.; Feig, M. Multi-state modeling of G-protein coupled receptors at experimental accuracy. Proteins: Struct., Funct., Bioinf. 2022, 90, 1873–1885.

(4) Bouvignies, G.; Vallurupalli, P.; Hansen, D. F.; Correia, B. E.; Lange, O.; Bah, A.; Vernon, R. M.; Dahlquist, F. W.; Baker, D.; Kay, L. E. Solution structure of a minor and transiently formed state of a T4 lysozyme mutant. Nature 2011, 477, 111–114.

(5) Bowman, G. R.; Geissler, P. L. Equilibrium fluctuations of a single folded protein reveal a multitude of potential cryptic allosteric sites. Proc. Natl. Acad. Sci. U. S. A. 2012, 109, 11681–11686.

(6) Csermely, P.; Palotai, R.; Nussinov, R. Induced fit, conformational selection and independent dynamic segments: an extended view of binding events. Trends Biochem. Sci. 2010, 35, 539–546.

(7) Saldaño, T.; Escobedo, N.; Marchetti, J.; Zea, D. J.; Mac Donagh, J.; Velez Rueda, A. J.; Gonik, E.; García Melani, A.; Novomisky Nechcoff, J.; Salas, M. N.; Peters, T.; Demitroff, N.; Fernandez Alberti, S.; Palopoli, N.; Fornasari, M. S.; Parisi, G. Impact of protein conformational diversity on AlphaFold predictions. Bioinformatics 2022, 38, 2742–2748.

(8) Del Alamo, D.; Sala, D.; Mchaourab, H. S.; Meiler, J. Sampling alternative conformational states of transporters and receptors with AlphaFold2. eLife 2022, 11, e75751.

(9) Abramson, J.; Adler, J.; Dunger, J.; Evans, R.; Green, T.; Pritzel, A.; Ronneberger, O.; Willmore, L.; Ballard, A. J.; Bambrick, J.; Bodenstein, S. W.; Evans, D. A.; Hung, C. C.; O’Neill, M.; Reiman, D.; Tunyasuvunakool, K.; Wu, Z.; Žemgulytė, A.; Arvaniti, E.; Beattie, C.; Bertolli, O.; Bridgland, A.; Cherepanov, A.; Congreve, M.; Cowen-Rivers, A. I.; Cowie, A.; Figurnov, M.; Fuchs, F. B.; Gladman, H.; Jain, R.; Khan, Y. A.; Low, C. M. R.; Perlin, K.; Potapenko, A.; Savy, P.; Singh, S.; Stecula, A.; Thillaisundaram, A.; Tong, C.; Yakneen, S.; Zhong, E. D.; Zielinski, M.; Žídek, A.; Bapst, V.; Kohli, P.; Jaderberg, M.; Hassabis, D.; Jumper, J. M. Accurate structure prediction of biomolecular interactions with AlphaFold 3. Nature 2024, 630, 493–500.

(10) Gopalan, S.; Narayanan, S. Hallucinations in AlphaFold3 for Intrinsically Disordered Proteins with disorder in Biological Process Residues. arXiv 2025, DOI: 10.48550/arXiv.2510.15939.

(11) Fang, M.; Wang, C.; Shi, J.; Lian, F.; Jin, Q.; Wang, Z.; Zhang, Y.; Chen, P.; Cui, Z.; Wang, Y.; Zhang, Z.; Ke, Y.; Han, Q.; Cao, L. HalluDesign: Protein Optimization and de novo Design via Iterative Structure Hallucination and Sequence design. bioRxiv 2025, DOI: 10.1101/2025.11.08.686881.

(12) Chai Discovery; Boitreaud, J.; Dent, J.; McPartlon, M.; Meier, J.; Reis, V.; Rogozhnikov, A.; Wu, K. Chai-1: Decoding the molecular interactions of life. bioRxiv 2024, DOI: 10.1101/2024.10.10.615955.

(13) Wohlwend, J.; Corso, G.; Passaro, S.; Getz, N.; Reveiz, M.; Leidal, K.; Swiderski, W.; Atkinson, L.; Portnoi, T.; Chinn, I.; Silterra, J.; Jaakkola, T.; Barzilay, R. Boltz-1: Democratizing biomolecular interaction modeling. bioRxiv 2024, DOI: 10.1101/2024.11.19.624167.

(14) Passaro, S.; Corso, G.; Wohlwend, J.; Reveiz, M.; Thaler, S.; Somnath, V. R.; Getz, N.; Portnoi, T.; Roy, J.; Stark, H.; Kwabi-Addo, D.; Beaini, D.; Jaakkola, T.; Barzilay, R. Boltz-2: Towards Accurate and Efficient Binding Affinity Prediction. bioRxiv 2025, DOI: 10.1101/2025.06.14.659707.

(15) Lewis, S.; Hempel, T.; Jiménez-Luna, J.; Gastegger, M.; Xie, Y.; Foong, A. Y. K.; Satorras, V. G.; Abdin, O.; Veeling, B. S.; Zaporozhets, I.; Chen, Y.; Yang, S.; Foster, A. E.; Schneuing, A.; Nigam, J.; Barbero, F.; Stimper, V.; Campbell, A.; Yim, J.; Lienen, M.; Shi, Y.; Zheng, S.; Schulz, H.; Munir, U.; Sordillo, R.; Tomioka, R.; Clementi, C.; Noé, F. Scalable emulation of protein equilibrium ensembles with generative deep learning. Science 2025, 389, eadv9817.

(16) Wang, Y.; Li, Y.; Chen, J.; Lai, L. Modeling protein–ligand interactions for drug discovery in the era of deep learning. Chem. Soc. Rev. 2025, 54, 11141–11183.

(17) Menon, K. M.; Davasam, A.; Chen, G.; Bryant, C.; Lam, B.; Yang, C.; Barcelos, D.; Liu, Y.; Liu, F.; Alon, A.; Lyu, J. AlphaFold3 for Structure-Guided Ligand Discovery. bioRxiv 2025, DOI: 10.64898/2025.12.04.692352.

(18) Kim, J.; Correy, G. J.; Hall, B. W.; Rachman, M. M.; Mailhot, O.; Togo, T.; Gonciarz, R. L.; Jaishankar, P.; Neitz, R. J.; Hantz, E. R.; Doruk, Y. U.; Stevens, M. G. V.; Diolaiti, M. E.; Reid, R.; Gopalkrishnan, S.; Krogan, N. J.; Renslo, A. R.; Ashworth, A.; Shoichet, B. K.; Fraser, J. S. Large scale prospective evaluation of co-folding across 557 Mac1-ligand complexes and three virtual screens. Elife 2026, 15, RP110475.

(19) Krokidis, M. G.; Koumadorakis, D. E.; Lazaros, K.; Ivantsik, O.; Exarchos, T. P.; Vrahatis, A. G.; Kotsiantis, S.; Vlamos, P. AlphaFold3: an overview of applications and performance insights. Int. J. Mol. Sci. 2025, 26, 3671.

(20) Chakravarty, D.; Schafer, J. W.; Chen, E. A.; Thole, J. F.; Ronish, L. A.; Lee, M.; Porter, L. L. AlphaFold predictions of fold-switched conformations are driven by structure memorization. Nat. Commun. 2024, 15, 7296.

(21) Chakravarty, D.; Lee, M.; Porter, L. L. Proteins with alternative folds reveal blind spots in AlphaFold-based protein structure prediction. Curr. Opin. Struct. Biol. 2025, 90, 102973.

(22) Jing, B.; Berger, B.; Jaakkola, T. AI-based Methods for Simulating, Sampling, and Predicting Protein Ensembles. Curr. Opin. Struct. Biol. 2026, 98, 103251.

(23) Sledzieski, S.; Hanson, S. RocketSHP: Ultra-fast proteome-scale prediction of protein dynamics. bioRxiv 2025, DOI: 10.1101/2025.06.12.659353.

(24) Prud, M.; Kyrychenko, A. Studying Ligand-Protein Interactions in the Era of Artificial Intelligence: Benchmarking Boltz-1 for 3D-Structure Prediction of Biomolecular Complexes. Kharkov Univ. Bull. Chem. Ser. 2025, 44, 57–71.

(25) Tanaka, Y.; Hipolito, C. J.; Maturana, A. D.; Ito, K.; Kuroda, T.; Higuchi, T.; Katoh, T.; Kato, H. E.; Hattori, M.; Kumazaki, K.; Tsukazaki, T.; Ishitani, R.; Suga, H.; Nureki, O. Structural basis for the drug extrusion mechanism by a MATE multidrug transporter. Nature 2013, 496, 247–251.

(26) Oh, B.-H.; Pandit, J.; Kang, C.-H.; Nikaido, K.; Gokcen, S.; Ames, G. F.; Kim, S.H. Three-dimensional structures of the periplasmic lysine/arginine/ornithine-binding protein with and without a ligand. J. Biol. Chem. 1993, 268, 11348–11355.

(27) Oh, B. H.; Kang, C. H.; De Bondt, H.; Kim, S. H.; Nikaido, K.; Joshi, A. K.; Ames, G. F. The bacterial periplasmic histidine-binding protein: structure/function analysis of the ligand-binding site and comparison with related proteins. J. Biol. Chem. 1994, 269, 4135–4143.

(28) Hunt, J. F.; Weinkauf, S.; Henry, L.; Fak, J. J.; McNicholas, P.; Oliver, D. B.; Deisenhofer, J. Nucleotide control of interdomain interactions in the conformational reaction cycle of SecA. Science 2002, 297, 2018–2026.

(29) Zimmer, J.; Nam, Y.; Rapoport, T. A. Structure of a complex of the ATPase SecA and the protein-translocation channel. Nature 2008, 455, 936–943.

(30) Dror, R. O.; Arlow, D. H.; Borhani, D. W.; Jensen, M. Ø.; Piana, S.; Shaw, D. E. Identification of two distinct inactive conformations of the β_2_ adrenergic receptor reconciles structural and biochemical observations. Proc. Natl. Acad. Sci. U. S. A. 2009, 106, 4689–4694.

(31) Kuroda, T.; Tsuchiya, T. Multidrug efflux transporters in the MATE family. Biochim. Biophys. Acta 2009, 1794, 763–768.

(32) Omote, H.; Hiasa, M.; Matsumoto, T.; Otsuka, M.; Moriyama, Y. The MATE proteins as fundamental transporters of metabolic and xenobiotic organic cations. Trends Pharmacol. Sci. 2006, 27, 587–593.

(33) Lu, M.; Radchenko, M.; Symersky, J.; Nie, R.; Guo, Y. Structural insights into H+-coupled multidrug extrusion by a MATE transporter. Nat. Struct. Mol. Biol. 2013, 20, 1310–1317.

(34) Zakrzewska, S.; Mehdipour, A. R.; Malviya, V. N.; Nonaka, T.; Koepke, J.; Muenke, C.; Hausner, W.; Hummer, G.; Safarian, S.; Michel, H. Inward-facing conformation of a multidrug resistance MATE family transporter. Proc. Natl. Acad. Sci. U. S. A. 2019, 116, 12275–12284.

(35) Oh, B.-H.; Ames, G. F.; Kim, S.-H. Structural basis for multiple ligand specificity of the periplasmic lysine-, arginine-, ornithine-binding protein. J. Biol. Chem. 1994, 269, 26323–26330.

(36) Felder, C. B.; Graul, R. C.; Lee, A. Y.; Merkle, H.-P.; Sadee, W. The Venus flytrap of periplasmic binding proteins: an ancient protein module present in multiple drug receptors. AAPS PharmSci. 1999, 1, E2.

(37) Chatzi, K. E.; Sardis, M. F.; Economou, A.; Karamanou, S. SecA-mediated targeting and translocation of secretory proteins. Biochim. Biophys. Acta - Mol. Cell Res. 2014, 1843, 1466–1474.

(38) Tsirigotaki, A.; De Geyter, J.; Šoštaric, N.; Economou, A.; Karamanou, S. Protein export through the bacterial Sec pathway. Nat. Rev. Microbiol. 2017, 15, 21–36.

(39) Bauer, B. W.; Rapoport, T. A. Mapping polypeptide interactions of the SecA ATPase during translocation. Proc. Natl. Acad. Sci. U. S. A. 2009, 106, 20800–20805.

(40) Bauer, B. W.; Shemesh, T.; Chen, Y.; Rapoport, T. A. A “push and slide” mechanism allows sequence-insensitive translocation of secretory proteins by the SecA ATPase. Cell 2014, 157, 1416–1429.

(41) Chen, Y.; Bauer, B. W.; Rapoport, T. A.; Gumbart, J. C. Conformational changes of the clamp of the protein translocation ATPase SecA. J. Mol. Biol. 2015, 427, 2348–2359.

(42) Ernst, I.; Haase, M.; Ernst, S.; Yuan, S.; Kuhn, A.; Leptihn, S. Large conformational changes of a highly dynamic pre-protein binding domain in SecA. Commun. Biol. 2018, 1, 130.

(43) Osborne, A. R.; Clemons Jr, W. M.; Rapoport, T. A. A large conformational change of the translocation ATPase SecA. Proc. Natl. Acad. Sci. U. S. A. 2004, 101, 10937–10942.

(44) Ou, X.; Ma, C.; Sun, D.; Xu, J.; Wang, Y.; Wu, X.; Wang, D.; Yang, S.; Gao, N.; Song, C.; Li, L. SecY translocon chaperones protein folding during membrane protein insertion. Cell 2025, 188, 1912–1924.e13.

(45) Motiejunaite, J.; Amar, L.; Vidal-Petiot, E. Adrenergic receptors and cardiovascular effects of catecholamines. Ann. Endocrinol. (Paris). 2021, 82, 193–197.

(46) Mersmann, H. J. β-adrenergic receptor modulation of adipocyte metabolism and growth. J. Anim. Sci. 2002, 80, E24–E29.

(47) Bang, I.; Choi, H.-J. Structural features of β2 adrenergic receptor: Crystal structures and beyond. Mol. Cells 2015, 38, 105–111.

(48) Rasmussen, S. G. F.; DeVree, B. T.; Zou, Y.; Kruse, A. C.; Chung, K. Y.; Kobilka, T. S.; Thian, F. S.; Chae, P. S.; Pardon, E.; Calinski, D.; Mathiesen, J. M.; Shah, S. T.; Lyons, J. A.; Caffrey, M.; Gellman, S. H.; Steyaert, J.; Skiniotis, G.; Weis, W. I.; Sunahara, R. K.; Kobilka, B. K. Crystal structure of the β_2_ adrenergic receptor–Gs protein complex. Nature 2011, 477, 549–555.

(49) Lamichhane, R.; Liu, J. J.; Pljevaljcic, G.; White, K. L.; van der Schans, E.; Ka-tritch, V.; Stevens, R. C.; Wüthrich, K.; Millar, D. P. Single-molecule view of basal activity and activation mechanisms of the G protein-coupled receptor β_2_AR. Proc. Natl. Acad. Sci. U. S. A. 2015, 112, 14254–14259.

(50) Nygaard, R.; Zou, Y.; Dror, R. O.; Mildorf, T. J.; Arlow, D. H.; Manglik, A.; Pan, A. C.; Liu, C. W.; Fung, J. J.; Bokoch, M. P.; Thian, F. S.; Kobilka, T. S.; Shaw, D. E.; Mueller, L.; Prosser, R. S.; Kobilka, B. K. The dynamic process of β_2_-adrenergic receptor activation. Cell 2013, 152, 532–542.

(51) Wayment-Steele, H. K.; Ojoawo, A.; Otten, R.; Apitz, J. M.; Pitsawong, W.; Hömberger, M.; Ovchinnikov, S.; Colwell, L.; Kern, D. Predicting multiple conformations via sequence clustering and AlphaFold2. Nature 2024, 625, 832–839.

(52) Mitjavila-Domenèch, N.; Díaz-Holguín, A.; Hu, H.; Kahlous, N. A.; Cabeza de Vaca, I.; Wallner, B.; Carlsson, J. Improving AlphaFold2 performance in virtual screens targeting GPCRs by enhancing binding-site conformational sampling. J. Chem. Inf. Model. 2026, In press.

(53) Stein, R. A.; Mchaourab, H. S. SPEACH AF: Sampling protein ensembles and conformational heterogeneity with AlphaFold2. PLoS Comput. Biol. 2022, 18, e1010483.

(54) Mirdita, M.; Schütze, K.; Moriwaki, Y.; Heo, L.; Ovchinnikov, S.; Steinegger, M. ColabFold: making protein folding accessible to all. Nat. Methods 2022, 19, 679–682.

(55) Bryant, P.; Noé, F. Structure prediction of alternative protein conformations. Nat. Commun. 2024, 15, 7328.

(56) Vani, B. P.; Aranganathan, A.; Wang, D.; Tiwary, P. AlphaFold2-RAVE: From sequence to Boltzmann ranking. J. Chem. Theory Comput. 2023, 19, 4351–4354.

(57) Gu, X.; Aranganathan, A.; Tiwary, P. Empowering AlphaFold2 for protein conformation selective drug discovery with AlphaFold2-RAVE. Elife 2024, 13, RP99702.

(58) Pang, Y. T.; Yang, L.; Kuo, K. M.; Gumbart, J. C. DeepPath: overcoming data scarcity for protein transition pathway prediction using physics-based deep learning. Chem. Sci. 2026, 17, 12055–12073.

(59) Agarwal, V.; McShan, A. C. The power and pitfalls of AlphaFold2 for structure prediction beyond rigid globular proteins. Nat. Chem. Biol. 2024, 20, 950–959.

(60) Kryshtafovych, A.; Schwede, T.; Topf, M.; Fidelis, K.; Moult, J. Critical assessment of methods of protein structure prediction (CASP)—Round XV. Proteins: Struct., Funct., Bioinf. 2023, 91, 1539–1549.

(61) Dube, N.; Ramelot, T. A.; Benavides, T. L.; Huang, Y. J.; Moult, J.; Kryshtafovych, A.; Montelione, G. T. Modeling alternative conformational states in CASP16. Proteins: Struct., Funct., Bioinf. 2026, 94, 330–347.

(62) Kryshtafovych, A.; Schwede, T.; Topf, M.; Fidelis, K.; Moult, J. Progress and bottlenecks for deep learning in computational structure biology: CASP Round XVI. Proteins: Struct., Funct., Bioinf. 2026, 94, 5–14.

(63) Stratiichuk, R.; Kyrylenko, R.; Koleiev, I.; Savchenko, I.; Voitsitskyi, T.; Husak, V.; Yesylevskyy, S.; Starosyla, S.; Nafiiev, A. Sampling and Ranking of Protein Conformations Using Machine Learning Techniques Do Not Improve the Quality of Rigid Protein–Protein Docking. J. Chem. Inf. Model. 2025, 65, 10167–10179.

(64) Abriata, L. A. The latest AI breakthroughs in structural biology: protein binder design and conformational state prediction. Commun. Biol. 2026, 9, 627.

(65) Sala, D.; Engelberger, F.; Mchaourab, H. S.; Meiler, J. Modeling conformational states of proteins with AlphaFold. Curr. Opin. Struct. Biol. 2023, 81, 102645.

(66) He, X.-h.; Li, J.-r.; Shen, S.-y.; Xu, H. E. AlphaFold3 versus experimental structures: assessment of the accuracy in ligand-bound G protein-coupled receptors. Acta Pharmacol. Sin. 2025, 46, 1111–1122.

(67) Zha, J.; Li, N.; Li, M.; Liu, X.; Zhu, R.; Feng, L.; Lu, X.; Zhang, J. Assessing the Performance of BioEmu in Understanding Protein Dynamics. Int. J. Mol. Sci. 2026, 27, 2896.

(68) Bhakat, S.; Strauch, E.-M. Accelerated Sampling of Protein Dynamics Using BioEmu-Augmented Molecular Simulation. J. Chem. Inf. Model. 2026, 66, 7168–7178.

(69) Yu, H.; Bekar-Cesaretli, A. A.; Lazou, M.; Kozakov, D.; Joseph-McCarthy, D.; Vajda, S. Bias in the AlphaFold3 prediction of ligand-induced domain motion in enzymes. Proc. Natl. Acad. Sci. U. S. A. 2026, 123, e2530709123.

(70) Shen, S.-y.; Li, J.-r.; Wang, Y.-s.; Li, S.n.; Xu, H. E.; He, X.-h. An update for AlphaFold3 versus experimental structures: assessing the precision of small molecule binding in GPCRs. Acta Pharmacol. Sin. 2025, 46, 3355–3364.

(71) Shen, C.; Zhang, X.; Gu, S.; Zhang, O.; Wang, Q.; Du, G.; Zhao, Y.; Jiang, L.; Pan, P.; Kang, Y.; Zhao, Q.; Hsieh, C.-Y.; Hou, T. Unlocking the application potential of AlphaFold3-like approaches in virtual screening. Chem. Sci. 2026, 17, 2858–2879.

(72) Zhang, L.; Friesner, R. A.; Miller, E. B.; Rodrigues, J. P. Generalization and Usability of Co-Folded GPCR–Ligand Complexes: A Physics-Guided Assessment. ChemRxiv 2026, DOI: 10.26434/chemrxiv-2026-1rkqz.

(73) The UniProt Consortium. UniProt: the universal protein knowledgebase in 2025. Nucleic Acids Res. 2025, 53, D609–D617.

(74) Berman, H. M.; Westbrook, J.; Feng, Z.; Gilliland, G.; Bhat, T. N.; Weissig, H.; Shindyalov, I. N.; Bourne, P. E. The Protein Data Bank. Nucleic Acids Res. 2000, 28, 235–242.

(75) Wayment-Steele, H. K.; Ovchinnikov, S.; Colwell, L.; Kern, D. Does Sequence Clustering Confound AlphaFold2? J. Mol. Biol. 2025, 437, 169376.

(76) PyMOL The PyMOL Molecular Graphics System, Version 3.1.6.1, Schrödinger, LLC.

(77) Zhang, Y.; Skolnick, J. TM-align: a protein structure alignment algorithm based on the TM-score. Nucleic Acids Res. 2005, 33, 2302–2309.

(78) Xu, J.; Zhang, Y. How significant is a protein structure similarity with TM-score = 0.5? Bioinformatics 2010, 26, 889–895.

