## Supporting Information for "Benchmarking AI Protein Structure Predictors Reveals a Persistent Bias in Multi-State Proteins"

Muhui Ye,[†] Yu-Hong Wang,[‡] Maximilian Brogi,[†] Jerry M. Parks,[¶] Katie M. Kuo,[*,§] and James C. Gumbart[*,§,‡]

†*School of Biological Sciences, Georgia Institute of Technology, Atlanta, GA, 30332 USA*

‡*School of Chemistry & Biochemistry, Georgia Institute of Technology, Atlanta, GA, 30332 USA*

¶*Biosciences Division, Oak Ridge National Laboratory, Oak Ridge, TN, 37831 USA*

§*School of Physics, Georgia Institute of Technology, Atlanta, GA, 30332 USA*

*

### Supporting Information

This document provides extended methods, additional analyses, and supplementary figures/tables referenced in the main text.

#### S1. Selecting conformations and conditions

To assess the ability of current structure predictors to capture functionally relevant conformational diversity, we selected four proteins with experimentally characterized conformational transitions: PfMATE, LAO, SecA, and $\beta_2$AR. These proteins represent diverse conformational mechanisms including alternating access transport, ligand-induced domain closure, ATP-dependent motor dynamics, and GPCR activation.

Protein sequences and annotations were obtained from the UniProt database.[1] Reference experimental structures were retrieved from the Protein Data Bank.[2] For PfMATE (Uniprot: Q8U2X0), only apo predictions were generated as both conformational states are accessible without ligands. For LAO (P02911), two conditions were tested: apo without ligands and holo with L-arginine specified via SMILES string. For SecA (P28366), three conditions were tested: apo without binding partners, ATP-bound, and in complex with the SecYEG translocon (SecY: Q9X1I9, SecE: P35874, SecG: P0AG90) with ADP. For $\beta_2$AR (P07550), four conditions were tested to span the activation pathway: apo; agonist (noradrenaline); agonist and G protein (G$\alpha$: P63092, G$\beta$: P62873, G$\gamma$: P59768); and agonist, G protein, and GTP. Small molecule ligands were specified using canonical SMILES 1.5.0 strings as input for the predictors. Protein binding partners were provided through full-length sequences. No distance constraints were applied. The counts of each conformation's representation in the PDB are provided in Table S1.

**Table S1: Distribution of experimentally determined structures across conformational states for the benchmark proteins studied.**

| Protein | State / Conformation | Count | PDB IDs | Total |
|---|---|---|---|---|
| PfMATE | Outward-facing | 10 | 3VVN, 3VVO, 3VVP, 3VVR, 3VVS, 3W4T, 3WBN, 4MLB, 6GWH, 6HFB | 11 |
| | Inward-facing | 1 | 6FHZ | |
| LAO[a] | Apo | 5 | 2LAO, 6MKX, 6ML0, 6MLD, 6MLV | 22 |
| | Holo | 17 | 1LST, 1LAF, 1LAG, 1LAH, 5OWF, 6FT2, 6MKU, 6MKW, 6ML9, 6MLA, 6MLE, 6MLG, 6MLI, 6MLJ, 6MLN, 6MLO, 6MLP | |
| SecA | Wide-open | 2 | 1M6N, 1M74 | 12 |
| | Open | 2 | 1TF2, 1TF5 | |
| | Closed | 8 | 3DIN, 3DL8, 8Y9Y, 8Y9Z, 8YA0, 8YA2, 8YA3, 8YAS | |
| $\beta_2$AR | Apo (inactive) | 19 | 2R4R, 2R4S, 2RH1, 3D4S, 3NY8, 3NY9, 3NYA, 5D5B, 5D6L, 5JQH, 5X7D, 6OBA, 6PS0, 6PS1, 6PS6, 8JJO, 8W1V, 9CHV, 9CHX | 126 |
| | Agonist-bound (inactive) | 17 | 3KJ6, 3P0G, 3PDS, 4GBR, 4LDL, 4LDO, 5D5A, 6PRZ, 6PS2, 6PS3, 6PS4, 6PS5, 8JJ8, 8JJL, 9CHU, 7XK9, 7XKA | |
| | Agonist + $G\alpha\beta\gamma$ (active) | 29 | 3SN6, 4LDE, 4QKX, 6MXT, 6N48, 6NI3, 7BZ2, 7DHI, 7DHR, 8GDZ, 8GE1, 8GE2, 8GE3, 8GE4, 8GE5, 8GE6, 8GE7, 8GE8, 8GE9, 8GEA, 8GEB, 8GEC, 8GED, 8GEE, 8GEF, 8GEG, 8GEH, 8GEI, 8GEJ | |
| | Agonist + $G\alpha\beta\gamma$ + GTP (active) | 61 | 6E67, 8GFX, 8GFV, 8GFW, 8GFY, 8GFZ, 8GG0, 8GG1, 8GG2, 8GG3, 8GG4, 8GG5, 8GG6, 8GG7, 8GG8, 8GG9, 8GGA, 8GGB, 8GGC, 8GGE, 8GGF, 8GGI, 8GGJ, 8GGK, 8GGL, 8GGM, 8GGN, 8GGO, 8GGP, 8GGQ, 8GGR, 8GGS, 8GGT, 8GGU, 8GGV, 8GGW, 8GGX, 8GGY, 8GGZ, 8GH0, 8GH1, 8UNL, 8UNM, 8UNN, 8UNO, 8UNP, 8UNQ, 8UNR, 8UNS, 8UNT, 8UNU, 8UNV, 8UNW, 8UNX, 8UNY, 8UNZ, 8UO0, 8UO1, 8UO2, 8UO3, 8UO4 | |

[a]6XKS is missing a portion of the structure, so it is not possible to determine whether it represents an apo or holo conformation for LAO protein.

### S2. Running the predictors and comparison

Predictors were run on NVIDIA A100 GPUs in a high-performance computing environment. To fully assess de novo structure prediction capabilities, structural templates and MSAs were not used across all predictors. BioEmu does not accept user-specified ligands or binding partners and therefore was run only under apo conditions.

For PfMATE, LAO, and SecA, predictions from AlphaFold3, Chai-1, and Boltz-2 each comprised 10 independent random seeds with five samples per seed, yielding 50 structures per condition. BioEmu does not account for ligands or multimers and so was used to generate apo structures only, using default sampling parameters with a batch size of 30 samples per run for PfMATE, LAO, and SecA, and 20 samples per run for $\beta_2$AR.

AlphaFold3 was run with 3 recycles (`num_recycles`=3), representing the number of iterations through the Pairformer module for refinement. Chai-1 was run with three recycles (`num_recycles`=3), step scale of 1.638 for diffusion sampling, and one batch for parallel processing. Boltz-2 was run with three recycles (`num_recycles`=3) and one trunk sample (`num_trunk_samples`=1) for structure generation.

All reference structures were obtained from the PDB (Table S2) and processed to retain only protein chains corresponding to the target sequence, removing ligands, ions, water molecules, non-protein chains, crystallographic artifacts, non-native residues introduced for crystallographic stabilization, and any engineered or fusion sequences not present in the native protein. Root-mean-square deviation (RMSD) calculations were performed over C$\alpha$ atoms using PyMOL's[3] structure-based CEAlign algorithm. For each predicted structure, RMSD values to all relevant reference conformations were computed and visualized by plotting RMSD distances between predictions and two reference states, one on each axis.

To complement RMSD analysis and validate global fold preservation, TM-scores were calculated for all predictions using TM-align.[4] TM-score is a length-independent metric (range 0–1) where values $> 0.5$ indicate the same fold.[5] TM-scores exceeding 0.5 for all predictions, combined with variable RMSD values, indicate that conformational differences reflect

**Table S2: Reference structures used for conformational comparisons.**

| Protein | Conformational State | PDB ID(s) |
|---|---|---|
| PfMATE | Inward-facing | 6FHZ |
| | Outward-facing | 6GWH |
| LAO | Apo (open) | 2LAO |
| | Holo (closed) | 1LAF |
| SecA | "Closed" (SecYEG-engaged) | 8YAS |
| | Open | 1TF2 |
| | Wide-open | 1M74 |
| $\beta_2$AR | Inactive/Intermediate | 2RH1, 9CHU |
| | Active | 8GEG, 8GG8 |

domain-level rearrangements rather than global misfolding (Figures S1–S4 and Table S3).

### S3. AF-Cluster and MSA subsampling baseline

As a baseline comparison independent of the newer architectures evaluated above (AlphaFold3, Boltz-2, Chai-1, and BioEmu), we applied AF-Cluster[7] to all four target proteins. AF-Cluster uses AlphaFold2 as its underlying structure predictor, but differs in that conformational diversity is driven by MSA-level manipulation rather than by the model architecture or training data of the newer tools. Full-length query sequences were used to generate multiple sequence alignments via the ColabFold MSA server.[8] Each MSA was partitioned into sequence-similarity clusters using `ClusterMSA.py` from the AF-Cluster repository (DBSCAN clustering, default parameters), yielding 295 clusters for LAO, 147 for PfMATE, 142 for $\beta_2$AR, and 4 for SecA. The small number of clusters obtained for SecA reflects limited sequence diversity in its available MSA, with cluster sizes of only 4–5 sequences each. For LAO, PfMATE, and $\beta_2$AR, the ten largest clusters by sequence count were selected for structure prediction; all four clusters were used for SecA. As controls, ten uniformly randomly subsampled MSAs of depth 10 (U10) and ten of depth 100 (U100) were also generated for each protein using the same script, to test whether the evolutionary structure captured by clustering, as opposed to simple MSA depth reduction, drives any observed conformational

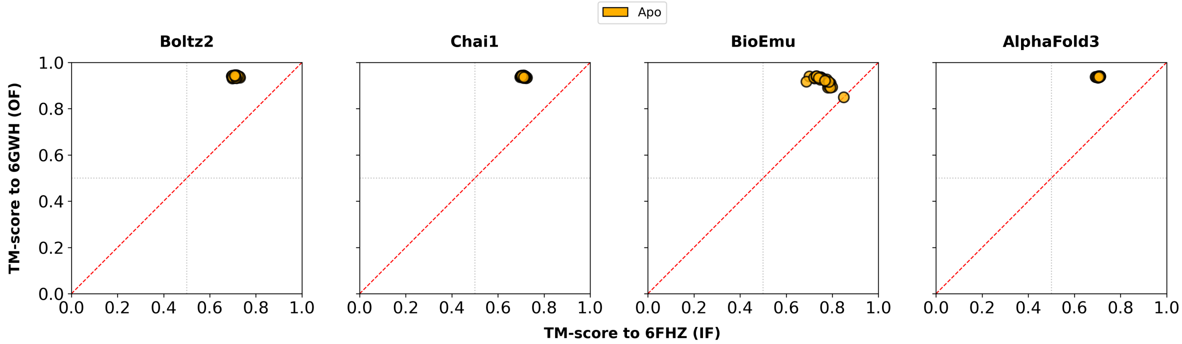


Figure S1: **TM-score analysis of PfMATE apo predictions.** Four-panel comparison of TM-score values for PfMATE predictions from Boltz-2, Chai-1, BioEmu, and AlphaFold3. Each point represents a predicted structure, with the x-axis showing TM-score to the inward-facing reference structure (6FHZ) and y-axis showing TM-score to the outward-facing reference structure (6GWH). The diagonal reference line (red dashed) represents equal similarity to both reference states; gray dotted lines indicate the TM-score = 0.5 threshold for same-fold classification.[5,6] Orange points represent apo predictions.

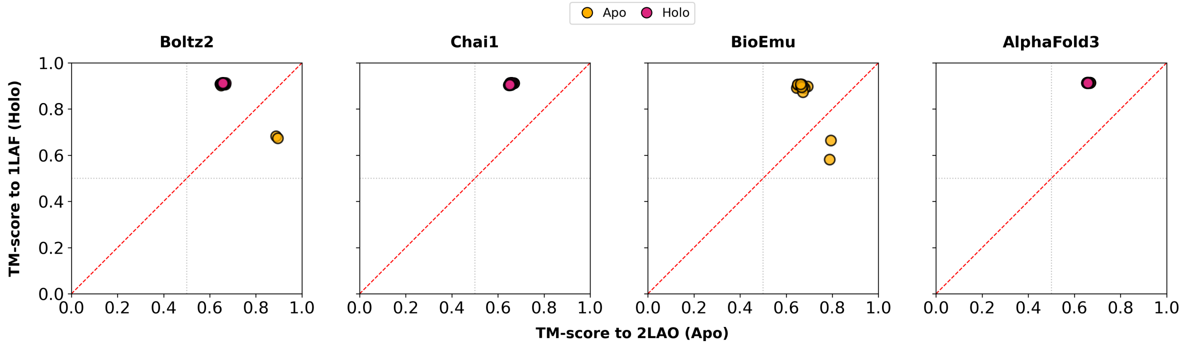


Figure S2: **TM-score analysis of LAO apo and holo predictions.** Four-panel comparison of TM-score values for LAO predictions from Boltz-2, Chai-1, BioEmu, and AlphaFold3. Each point represents a predicted structure, with the x-axis showing TM-score to the apo reference structure (2LAO) and the y-axis showing TM-score to the holo reference structure (1LAF). Orange points represent apo condition predictions; pink points represent holo condition predictions.

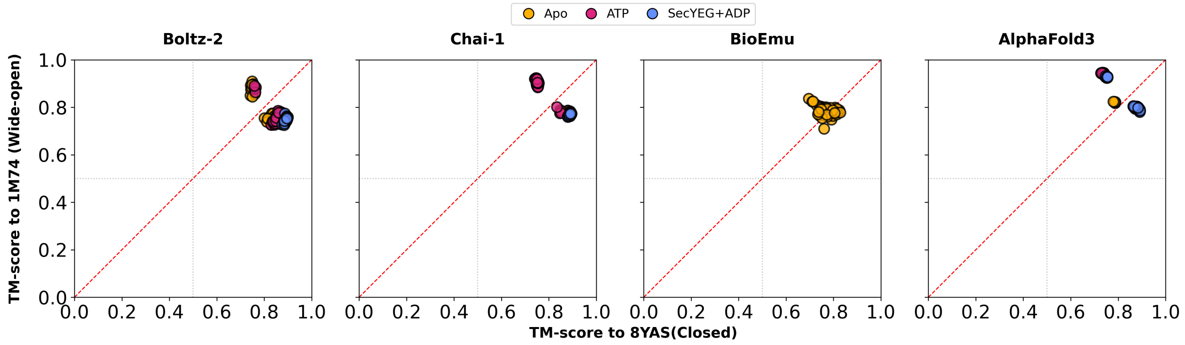


Figure S3: **TM-score analysis of SecA predictions under three conditions.** Four-panel comparison of TM-score values for SecA predictions from Boltz-2, Chai-1, BioEmu, and AlphaFold3. Each point represents a predicted structure, with the x-axis showing TM-score to the closed reference structure (8YAS) and the y-axis showing TM-score to the wide-open reference structure (1M74). Orange points represent apo predictions; pink points represent ATP-bound predictions; blue points represent SecYEG+ADP complex predictions.

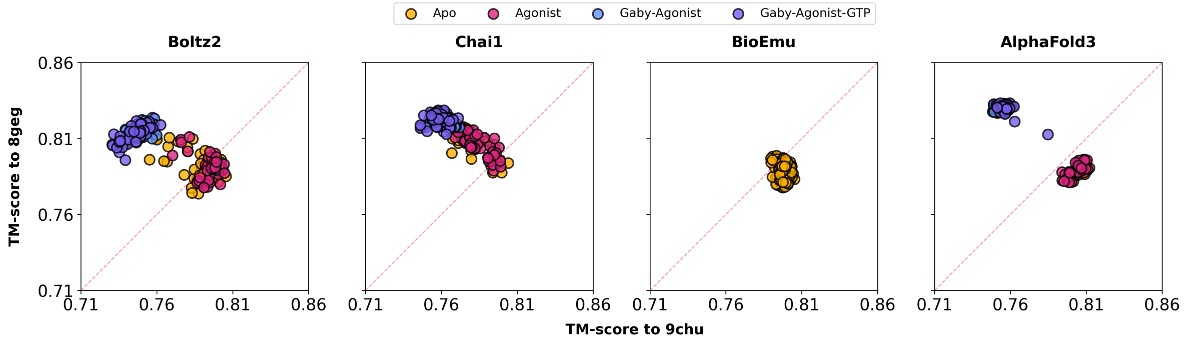


Figure S4: **TM-score analysis of $\beta_2$AR predictions under four conditions.** Four-panel comparison of TM-score values for $\beta_2$AR predictions from Boltz-2, Chai-1, BioEmu, and AlphaFold3. Each point represents a predicted structure, with the y-axis showing TM-score to the active reference structure (8GEG) and x-axis showing TM-score to the inactive reference structure (9CHU). Orange points represent apo predictions; pink points represent agonist-bound predictions; light blue points represent G protein-agonist complex predictions; teal points represent the full G protein-agonist-GTP condition.

**Table S3: Summary of TM-score statistics across all protein systems and prediction methods.**

| Protein | Method | Condition | TM to Ref 1 | TM to Ref 2 |
|---|---|---|---|---|
| *PfMATE (Ref 1: 6FHZ IF, Ref 2: 6GWH OF)* | | | | |
| | Boltz-2 | Apo | 0.695–0.730 | 0.931–0.945 |
| | Chai-1 | Apo | 0.696–0.725 | 0.932–0.944 |
| | BioEmu | Apo | 0.687–0.849 | 0.850–0.941 |
| | AlphaFold3 | Apo | 0.692–0.711 | 0.934–0.941 |
| *LAO (Ref 1: 2LAO Apo, Ref 2: 1LAF Holo)* | | | | |
| | Boltz-2 | Apo | 0.646–0.895 | 0.674–0.911 |
| | Boltz-2 | Holo | 0.651–0.668 | 0.911–0.913 |
| | Chai-1 | Apo | 0.648–0.669 | 0.905–0.913 |
| | Chai-1 | Holo | 0.647–0.661 | 0.904–0.913 |
| | BioEmu | Apo | 0.645–0.794 | 0.582–0.907 |
| | AlphaFold3 | Apo | 0.655–0.668 | 0.913–0.914 |
| | AlphaFold3 | Holo | 0.654–0.660 | 0.913–0.914 |
| *SecA (Ref 1: 8YAS Closed, Ref 2: 1M74 Wide-Open)* | | | | |
| | Boltz-2 | Apo | 0.742–0.870 | 0.726–0.909 |
| | Boltz-2 | ATP | 0.758–0.887 | 0.726–0.891 |
| | Boltz-2 | SecYEG+ADP | 0.875–0.899 | 0.726–0.774 |
| | Chai-1 | Apo | 0.741–0.880 | 0.775–0.923 |
| | Chai-1 | ATP | 0.742–0.889 | 0.776–0.923 |
| | Chai-1 | SecYEG+ADP | 0.880–0.895 | 0.759–0.780 |
| | BioEmu | Apo | 0.696–0.829 | 0.710–0.836 |
| | AlphaFold3 | Apo | 0.740–0.788 | 0.818–0.935 |
| | AlphaFold3 | ATP | 0.728–0.740 | 0.940–0.946 |
| | AlphaFold3 | SecYEG+ADP | 0.747–0.892 | 0.780–0.932 |
| *$\beta_2$AR (Ref 1: 9CHU Inactive, Ref 2: 8GEG Active)* | | | | |
| | Boltz-2 | Apo | 0.757–0.804 | 0.773–0.813 |
| | Boltz-2 | Agonist | 0.773–0.804 | 0.779–0.812 |
| | Boltz-2 | Gαβγ-Agonist | 0.737–0.760 | 0.807–0.825 |
| | Boltz-2 | Gαβγ-Agonist-GTP | 0.730–0.762 | 0.799–0.821 |
| | Chai-1 | Apo | 0.755–0.802 | 0.788–0.820 |
| | Chai-1 | Agonist | 0.768–0.801 | 0.786–0.818 |
| | Chai-1 | Gαβγ-Agonist | 0.746–0.771 | 0.817–0.827 |
| | Chai-1 | Gαβγ-Agonist-GTP | 0.748–0.770 | 0.814–0.827 |
| | BioEmu | Apo | 0.793–0.802 | 0.780–0.796 |
| | AlphaFold3 | Apo | 0.799–0.810 | 0.784–0.795 |
| | AlphaFold3 | Agonist | 0.795–0.809 | 0.781–0.794 |
| | AlphaFold3 | Gαβγ-Agonist | 0.750–0.757 | 0.829–0.831 |
| | AlphaFold3 | Gαβγ-Agonist-GTP | 0.752–0.782 | 0.815–0.831 |

shift.

Each selected sub-MSA was used as direct input to AlphaFold2 via `colabfold_batch`,[8] generating one structural model per sub-MSA (`num_models`=1, `num_recycle`=3, `model_type` =alphafold2, with no additional templates). RMSD calculations for AF-Cluster, U10, and U100 predictions were performed identically to those described in S2, using PyMOL's CEAlign algorithm against the same reference structures (Table S2). For SecA, AF-Cluster outputs were additionally evaluated using the PBD–HWD distance and PBD–hinge–NBD2 angle metrics described in Section S4.

#### S4. SecA domain-geometry state metrics

We quantified the conformational ensemble of each SecA prediction using two domain-level metrics. Domain boundaries were defined based on the P28366 (UniProt) sequence numbering: PBD (residues 216–357), HWD (residues 619–704), and NBD2 (residues 397–574).

(i) The PBD–HWD distance is defined as the Euclidean distance between the Cα centroids of the PBD and HWD domains.

(ii) The PBD–hinge–NBD2 angle is defined as the angle formed by three points: the Cα centroid of the PBD, the midpoint of the Cα atoms of residues 216 and 357 as the hinge vertex, and the Cα centroid of NBD2 .

These metrics were computed using PyMOL[3] for all predicted structures and reference experimental structures (1M74, 1TF2, 8YAS).

**Table S4: Summary of pTM and mean pLDDT across all protein systems and prediction methods. BioEmu does not output confidence scores by design.**

| Protein | Method | Condition | pTM | pLDDT |
|---|---|---|---|---|
| *PfMATE* | | | | |
| | AlphaFold3 | Apo | 0.928-0.930 | 88.0-89.1 |
| | Chai-1 | Apo | 0.936-0.938 | 88.5-88.7 |
| | Boltz-2 | Apo | 0.871-0.908 | 90.8-93.4 |
| | BioEmu | Apo | N/A | |
| *LAO* | | | | |
| | AlphaFold3 | Apo | 0.900-0.910 | 92.1-92.5 |
| | AlphaFold3 | Holo | 0.890-0.892 | 93.3-93.6 |
| | Chai-1 | Apo | 0.921-0.925 | 93.7-94.0 |
| | Chai-1 | Holo | 0.927-0.931 | 95.0-95.2 |
| | Boltz-2 | Apo | 0.729-0.856 | 89.9-91.1 |
| | Boltz-2 | Holo | 0.929-0.952 | 95.1-96.2 |
| | BioEmu | Apo | N/A | |
| *SecA* | | | | |
| | AlphaFold3 | Apo | 0.830-0.860 | 84.5-86.1 |
| | AlphaFold3 | ATP | 0.850-0.870 | 85.1-86.3 |
| | AlphaFold3 | SecYEG+ADP | 0.720-0.760 | 75.0-75.1 |
| | Chai-1 | Apo | 0.820-0.855 | 82.1-83.2 |
| | Chai-1 | ATP | 0.803-0.835 | 77.5-79.7 |
| | Chai-1 | SecYEG+ADP | 0.843-0.847 | 80.4-80.7 |
| | Boltz-2 | Apo | 0.671-0.752 | 76.2-79.5 |
| | Boltz-2 | ATP | 0.702-0.778 | 76.8-80.2 |
| | Boltz-2 | SecYEG+ADP | 0.791-0.877 | 72.3-75.2 |
| | BioEmu | Apo | N/A | |
| $\beta_2 AR$ | | | | |
| | AlphaFold3 | Apo | 0.710–0.720 | 70.2–71.9 |
| | AlphaFold3 | Agonist | 0.680–0.700 | 69.1–70.7 |
| | AlphaFold3 | Gαβγ-Agonist | 0.760–0.770 | 78.9–80.0 |
| | AlphaFold3 | Gαβγ-Agonist-GTP | 0.740–0.780 | 76.0–79.6 |
| | Chai-1 | Apo | 0.786–0.793 | 73.1–74.1 |
| | Chai-1 | Agonist | 0.781–0.789 | 72.4–73.2 |
| | Chai-1 | Gαβγ-Agonist | 0.810–0.822 | 80.4–80.9 |
| | Chai-1 | Gαβγ-Agonist-GTP | 0.807–0.816 | 80.4–80.9 |
| | Boltz-2 | Apo | 0.703–0.783 | 68.4–73.5 |
| | Boltz-2 | Agonist | 0.771–0.938 | 73.8–77.9 |
| | Boltz-2 | Gαβγ-Agonist | 0.681–0.879 | 72.0–74.9 |
| | Boltz-2 | Gαβγ-Agonist-GTP | 0.665–0.901 | 71.7–75.0 |
| | BioEmu | Apo | N/A | |

### References


(1) The UniProt Consortium. UniProt: the universal protein knowledgebase in 2025. *Nucleic Acids Res.* **2025**, *53*, D609–D617.

(2) Berman, H. M.; Westbrook, J.; Feng, Z.; Gilliland, G.; Bhat, T. N.; Weissig, H.; Shindyalov, I. N.; Bourne, P. E. The Protein Data Bank. *Nucleic Acids Res.* **2000**, *28*, 235–242.

(3) PyMOL The PyMOL Molecular Graphics System, Version 3.1.6.1, Schrödinger, LLC.

(4) Zhang, Y.; Skolnick, J. TM-align: a protein structure alignment algorithm based on the TM-score. *Nucleic Acids Res.* **2005**, *33*, 2302–2309.

(5) Xu, J.; Zhang, Y. How significant is a protein structure similarity with TM-score = 0.5? *Bioinformatics* **2010**, *26*, 889–895.

(6) Zhang, Y.; Skolnick, J. Scoring function for automated assessment of protein structure template quality. *Proteins* **2004**, *57*, 702–710.

(7) Wayment-Steele, H. K.; Ovchinnikov, S.; Colwell, L.; Kern, D. Does Sequence Clustering Confound AlphaFold2? *J. Mol. Biol.* **2025**, *437*, 169376.

(8) Mirdita, M.; Schütze, K.; Moriwaki, Y.; Heo, L.; Ovchinnikov, S.; Steinegger, M. ColabFold: making protein folding accessible to all. *Nat. Methods* **2022**, *19*, 679–682.
